# A look inside the net: freshwater turtles assort with conspecifics in feeding aggregations

**DOI:** 10.64898/2026.04.13.718235

**Authors:** Caitlin Menzies, Remus James, Julia Riley, Christina M. Davy, Roslyn Dakin

**Affiliations:** Department of Biology, Carleton University, 1125 Colonel By Drive, Ottawa, Ontario, Canada, K1S 5B6; Environmental and Life Science Program, Trent University, 1600 West Bank Drive, Peterborough, Ontario, Canada, K9L 0G2; Department of Biology, Mount Allison University, Flemington Building, 63B York St., Sackville, New Brunswick, Canada, E4L 1E2

**Keywords:** social evolution, social networks, behavioural ecology, sociality, freshwater turtles

## Abstract

Historically, non-avian reptiles were assumed to be non-social – yet recent studies show diverse social behaviours in squamates, crocodilians, and turtles. Here, we investigate social structure within feeding aggregations of three freshwater turtle species caught in baited traps in a coastal marsh over 12 years. In 488 instances in which traps contained turtles, 45% contained multiple individuals, and these aggregations were strongly positively assorted by species. Midland painted turtles (*Chrysemys picta marginata*) and Blanding’s turtles (*Emydoidea blandingii*) were captured with conspecifics more often than expected in non-social null models. Snapping turtles (*Chelydra serpentina*), the largest species in this study, were captured with conspecifics at a frequency similar to that predicted by null models, and were avoided by heterospecifics. This suggests that species differences play a role in how feeding aggregations are structured, with painted and Blanding’s turtles driving strong positive species assortment while avoiding snapping turtles around food sources. We did not detect significant intraspecific sex assortment in any of the three species, nor did turtle body size strongly affect the probability of aggregating with conspecifics at the perceived food source in the traps. Our study illustrates how long-term monitoring data can be used to investigate social structure in wild populations, an approach that may be valuable for species of conservation concern.

**Significance Statement:** Reptile sociality has been historically overlooked, but recent work has revealed intriguing social behaviours in non-avian reptiles. We investigated associations among three species of freshwater turtle, captured in baited traps over 12 years of trapping. Turtles in these feeding aggregations grouped with their own species more often than expected by chance. This result was mainly driven by the two smaller-bodied species (midland painted turtle and Blanding’s turtle), which were more likely to be caught with conspecifics than with other species. In addition, midland painted turtles and Blanding’s turtles showed avoidance of the larger snapping turtles. Our analyses illustrate how long-term monitoring data can be used to uncover previously unrecognized social behaviour in turtles and other species in which behaviour is difficult to observe directly.

## Introduction

Social behaviour is diverse across vertebrates and ranges from solitary living species, through varying levels of gregariousness, to the most extreme state of eusociality (Allaby 2020). Broadly, sociality can be defined as animals’ tendency to associate with other individuals (Gartland et al. 2021; Clark and Ioannou 2025) and can include the tendency to live in groups (Allaby 2020). Sociality is hypothesized to arise from initially non-social aggregations, where positive fitness outcomes arising from interactions among group members (such as reduced predation risk and increased reproductive and foraging success) eventually lead to the evolution of more complex social behaviours, like long-term kin-based social groups, altruism, and cooperation (Bode and Delcourt 2013; Philippe et al. 2016; Silva 2022). For example, feeding aggregations at localized resources are a common cause of resource-based clustering in the wild that may initially be driven by factors that are not inherently social in nature (Bode and Delcourt 2013; Wilson 2000).

Social network analyses can be used to explore how species, sex, physical condition, age, dominance, or other traits predict the structure of aggregations and social interactions (Kurvers et al. 2014; Krause et al. 2015; Webber and Vander Wal 2019; Rouleau et al. 2026). Social network analyses organize animal assemblages into a network of individuals, represented by nodes, which are associated in a predefined way (e.g., proximity, cooperation, relatedness, etc.), with the associations or “edges” representing connections between nodes (Croft et al. 2008). Nodes are often shown visually as points and lines connect these nodes and represent the edges. Within a social network, the assortment coefficient quantifies the extent to which similar individuals are associated with each other (Krause et al. 2015). In mixed-species communities, positive assortment by species indicates a tendency for individuals to associate with conspecifics. Within a species, positive assortment by sex indicates individuals’ tendency to associate with same-sex conspecifics (Newman 2003). Assortativity varies across and within species, depending on a variety of factors, which may include the distribution of environmental resources or their social and mating systems. For example, there is a plethora of evidence of birds joining mixed-species feeding aggregations (Schreffler et al. 2010; Latta and Wunderle 1996), and sex-based aggregations have been documented in marine iguanas and marine snakes (Wearmouth and Sims 2008).

Despite the shared ancestry of birds and turtles, the scientific community has historically considered turtles and other non-avian reptiles as solitary, non-social animals (Doody et al. 2013, 2021; Wilson 2000). Nevertheless, social behaviours documented in turtles include recognition of kin and familiar conspecifics (Ibáñez et al. 2013; Whitear et al. 2016; Versace et al. 2018), communal overwintering (Litzgus et al. 1999; Newton and Herman 2009; Rasmussen and Litzgus 2010; Bulté et al. 2024), social basking (Polo-Cavia et al. 2010; Russell 2016; Rouleau et al. 2026), communal nesting (Kell et al. 2022), vocal communication (Giles et al. 2009; Doody et al. 2013, 2021; Ferrara et al. 2013; Lacroix et al. 2022, 2025), social play (Kramer and Burghardt 1998; Burghardt 2005), early evidence of group living (McKnight et al 2023), and possible parental care (Ferrara et al. 2013; Topping and Valenzuela 2021). Understanding the full range of reptile social behaviour will help us understand the factors that drive the origin and persistence of social behaviour across vertebrates (Doody et al. 2013, 2021; Lacroix et al. 2025).

Basking aggregations are the best-studied example of grouping in freshwater turtles, likely because basking is a relatively conspicuous behaviour. Basking turtles expose all or part of their body to solar radiation while remaining immobile, either at the surface of the water or after climbing fully out of the water onto a raised object (Lindeman 1999; Bulté and Blouin-Demers 2010). Basking allows turtles to thermoregulate and shed parasitic leeches, and may also be used to induce behavioural fevers to fight infection (Zipko 1982; Lindeman 1999; Bulté and Blouin-Demers 2010; McKnight et al 2023; Chessman 2024). Basking aggregations are also thought to have a social function, facilitating increased vigilance against predators, hierarchy structures, and preferred associations (Zipko 1982; Lindeman 1999; Russel 2016; Lovich and Gibbons 2021; McKnight et al 2026; Rouleau et al 2026). Multi-species basking aggregations are common among Emydid turtles (Zipko 1982; Lovich and Gibbons 2021; Rouleau et al. 2026), with some evidence that larger turtles may displace their smaller counterparts (Lindeman 1999). In contrast, snapping turtles (*Chelydra serpentina*) rarely emerge to bask with conspecifics or other turtles (Obbard and Brooks 1979; Lovich and Gibbons 2021), notwithstanding isolated observations of a painted turtle using a snapping turtle carapace as a basking location (Legler 1956). Globally, interspecific variation is also reported in basking behaviours of other freshwater turtle species (e.g., Clavijo-Baquet and Magnone 2017; McKnight et al. 2023; Kidman et al. 2024).

However, most freshwater turtle behaviour occurs underwater, often in systems with high turbidity where direct observation is challenging. As a result, our understanding of aquatic social interactions in turtles lags behind our understanding of basking, and relies heavily on anecdotal observations. It is possible that aggregation underwater (for example, around food sources) might confer similar benefits to basking aggregations, for example, increased vigilance against predators such as North American river otters (*Lutra canadensis*). Observational reports reveal that painted turtles will enter hoop traps when snapping turtles are already present (Frazer et al. 1990; Ernst and Lovich 2009), but it is unclear how common this behaviour is. Painted turtles and snapping turtles also exhibit a cleaning symbiosis in which painted turtles remove algae and leeches from the carapaces of snapping turtles (Krawchuk et al. 1997). While observations of this behaviour are rare, it nevertheless illustrates how heterospecific social interactions can mutually benefit freshwater turtles.

In addition to species identity, interactions among turtles may also be influenced by body size (Luiselli et al. 2021). Among conspecifics, body size can affect competition for resources such as food or mates, with larger individuals tending to have the advantage (Lindeman 1999; Keevil et al. 2017), although the role of competition in shaping turtle communities remains uncertain (Luiselli 2008; Petrov et al. 2020). Some large turtles can also consume smaller turtles, so encounters with large individuals may increase a small turtle’s risk of injury or mortality (Polis and Myers 1985; Williams et al. 2026). Body size may also influence dominance hierarchies: in wood turtles (*Glyptemys insculpta*), larger individuals rank higher in intraspecific dominance hierarchies expressed during terrestrial mating behaviours (Kaufmann 1992). Experimental work shows that turtles can use chemosensory cues to recognize conspecifics and potential predators, and to infer the size and relative condition of potential mates or competitors (e.g., Ibáñez et al. 2012, 2014, 2015; Whitear et al. 2016). These abilities could enable turtles to assess the costs or benefits of aggregation at a localized resource, such as a perceived source of food. However, we are not aware of any studies that have compared social behaviours during feeding among co-occurring freshwater turtle species.

Here, we used social network analysis to investigate sociality in feeding aggregations of three freshwater turtle species in Ontario, Canada (Fig. 1A). We used data from a long-term monitoring project in which baited hoop traps captured Blanding’s turtles (*Emydoidea blandingii*), painted turtles (*Chrysemys picta*), and snapping turtles. These trapping activities within a continuous connected marsh habitat generated temporary aggregations around a perceived source of food. Although clustering at a resource does not require attraction to other individuals, a clustered distribution is a prerequisite for many types of social behaviour (Wilson 2000; Philippe et al. 2016; Allaby 2020). We consequently hypothesized that if foraging turtles assort with conspecifics, then turtles would be spatiotemporally clustered nonrandomly in the traps, and that aggregations in the traps would include more conspecific dyads than expected by chance, and fewer heterospecific dyads. We compared species assortment in networks generated from trapping data to expectations of a non-social null model to test our predictions. We then investigated a series of potential drivers of aggregation and assortment: (1) by testing whether co-occurrence frequencies within and between each species differed from non-social null expectations; (2) by examining the proportion of individuals in each focal species that co-occurred with at least one conspecific; and (3) by investigating whether body size variation within each species predicted co-capture. Finally, to investigate whether turtles might enter a trap to access potential mates, we tested for assortment by sex in the trap data for each species. Together, our analyses provide insight into the intra- and inter-specific interactions of these three species during underwater behaviours that are difficult to directly observe.

**Fig. 1.**
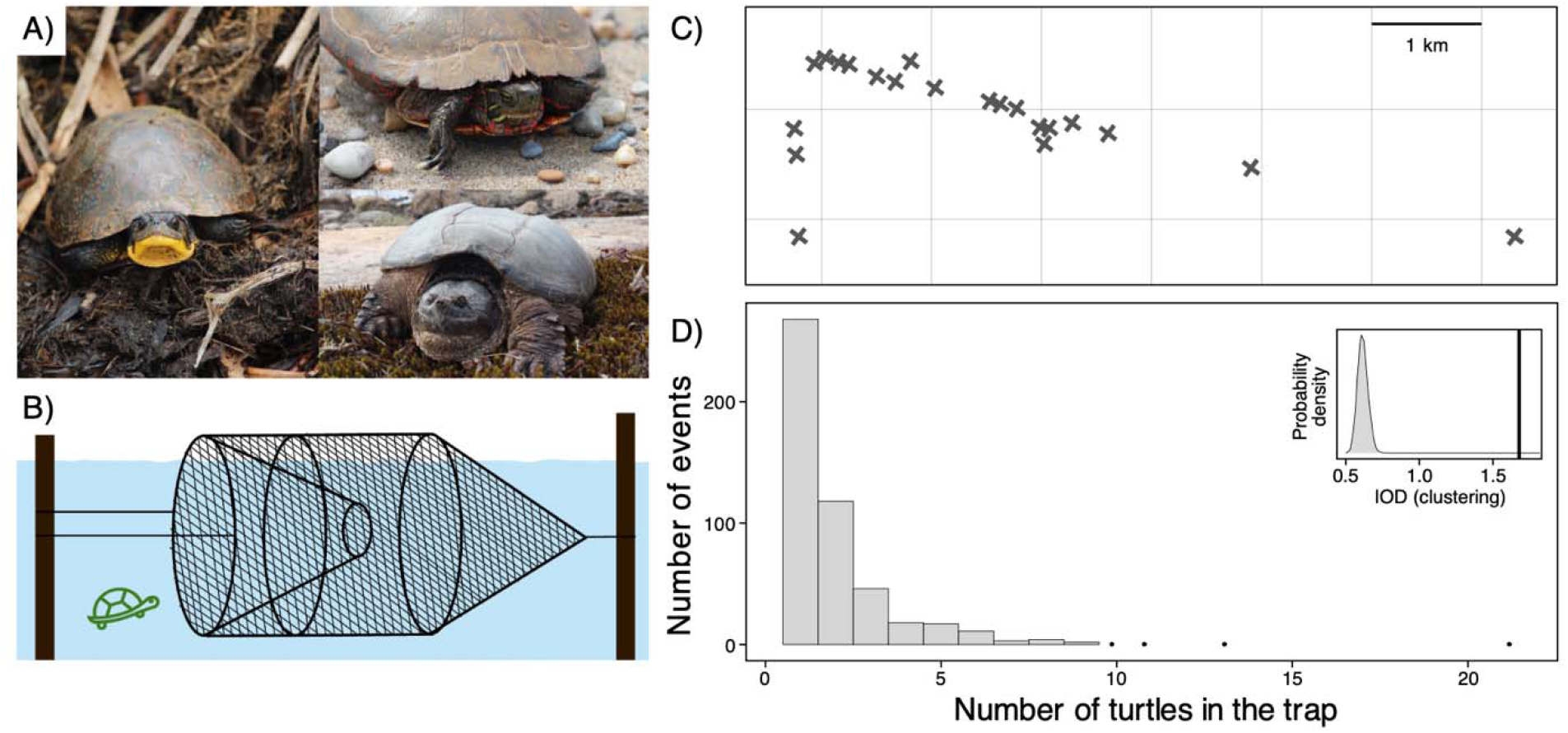
We studied associations between freshwater turtles that co-occurred in feeding aggregations captured in baited hoop traps. **(A)** This study included three species: the Blanding’s turtle (*Emydoidea blandingii*) (left), midland painted turtle (*Chrysemys picta marginata*) (top right), and snapping turtle (*Chelydra serpentina*) (bottom right). Photographs by Caitlin Menzies. **(B)** An illustration of a hoop trap. **(C)** Arrangement of 21 trap locations used in this study; note that not all locations were sampled in each year of the study. All locations are part of one contiguous marsh. **(D)** Distribution shown using a histogram of the number of turtles captured per capture event. For clarity, bars are omitted for x-values with only one event, and a point is shown for those x-values instead. The inset panel shows the index of dispersion (IOD) of 1.68 for these capture events (vertical line), with the null expectation for IOD shown by the grey probability density

## Methods

### Field methods

Data for this study were collected over 12 years (2009−2019 and 2022), as part of a long-term research and monitoring program in a coastal wetland on the north shore of Lake Erie, on the traditional territories of the Attiwonderonk and Missisauga nations. The wetland includes marshy areas dominated by cattails (*Typha* spp.), common reed (*Phragmites australis*), open meadow marsh, and coastal dunes. We set hoop traps (TN315 Turtle Nets, Memphis Net and Twine Co., Memphis, Tennessee; hoop diameter 91.4 cm; 3.8 cm^2^ mesh) in an 8.3 km^2^ region of the marsh that is used by all three study species. The trapping area contains sandy ridges separating shallow, marshy areas, with no physical barriers to turtle movement through the marsh (Fig. 1C). Traps were baited with sardines canned in oil (Fig. 1B), and were checked and rebaited at least twice daily, in the same sequence in which they had been baited, with approximately 12 hours overnight between checks. Average pairwise distance between hoop traps was 1.65 km (SD =1.87 km; max = 6.28 km; min = 0.13 km). It was not possible for a turtle released at one trapping site to enter a nearby trap before it was checked, although turtles could easily move among trap locations given more time.

Using trapping data from a long-term monitoring program for a behavioural study requires a few caveats, because the trapping methods were not designed with our current study in mind. Trapping dates and intensity varied substantially among years (Table S1), but our research questions focused on the composition of trapped aggregations of turtles, which we do not expect to be influenced by trap effort. The same trap types (i.e., the same hoop diameter and mesh size) were always used. Specific trap locations (i.e., GPS coordinates) varied among years as rising and falling lake levels altered the water depth across the coastal marsh. However, general trapping areas (i.e., particular shallow ponds within the marsh and the general locations of traps within those ponds) became more consistent over time, and concurrent visual and hand-capture surveys confirmed that the three target species were easily detected at all the trapping locations. Thus, we considered potential variation in the composition of turtle aggregations due to differences in habitat selection among the three species to be negligible.

For the present study, we defined a capture event as an instance in which a trap was checked and contained one or more turtles. Each captured turtle was marked by filing a unique pattern of notches into the marginal scutes (Cagle 1939). In some years, snapping turtles were also marked with a passive integrated transponder (PIT) tag (12 mm; Biomark, Boise, Idaho) that was inserted subcutaneously along the ventral surface of the hind leg. We measured body size by taking the curved carapace length (CCL) in mm with a flexible tape measure from the anterior edge of the nuchal scute to the posterior edge of the marginal scutes. We determined sex using a combination of species-specific features (Ernst and Lovich 2009; Lovich and Gibbons 2021). Specifically, Blanding’s turtles were sexed by examining the plastron (strongly concave in males, and flat or slightly convex in females). Painted turtles were sexed by examining the foreclaws (elongated in males and short in females), and tail morphology (short tail, cloaca located at or within the margin of the carapace on females; long, thicker tail with the cloaca located well beyond the margin of the carapace on males). Snapping turtles were sexed by comparing the length of the posterior lobe of the plastron to the distance from the base of the tail to the cloaca (posterior lobe longer than distance to cloaca in females, and shorter in males). It was not possible to record data blind because our study involved focal animals in the field.

### Data analysis

All data analyses were conducted using R 4.5.2 (R Core Team 2025) and ‘*tidyverse’* packages (Wickham et al. 2019). Our dataset included 491 capture events (instances in which a trap was checked and contained one or more turtles), in which 816 unique focal turtles were captured (128 Blanding’s turtles, 256 painted turtles, and 432 snapping turtles). A further seven individuals from other, non-focal turtle species were also captured, but are omitted from the analysis here due to their very low capture rates. Of the 816 individual turtles from the three focal species, 615 were captured with one or more conspecifics. Additionally, 115 focal individuals were captured more than once. We were able to confidently determine focal individuals’ sex in ∼92% of captures. Some smaller individuals were not yet exhibiting clear sexual dimorphism. These fell in the 8-9^th^ percentile of their species’ CCL size distributions, and we categorized the sex of these individuals as indeterminate.

### Clustering in feeding aggregations

We first investigated whether turtles in a capture event were more clustered than expected by random chance. To do this, we used the index of dispersion (IOD), computed as the variance of the number of turtles per capture event (i.e., an instance in which the trap was checked and contained at least one turtle), divided by the mean number of turtles per capture event (Perry and Hewitt 1991; Clark and Ioannou 2025). Larger values of IOD indicate a more aggregated distribution (Perry and Hewitt 1991; Clarke et al. 2006; Lester and McVinish 2016; Clark and Ioannou 2025). We compared the observed IOD to a null expectation for IOD that retained the same number of capture events and individual turtles per year, but randomly assigned individual turtles to capture events independently of each other. We calculated the expected IOD for 10,000 instances of this null simulation. Because instances in which traps were checked but empty (0s) were not consistently recorded, our null expectation for the IOD based on capture events differs from a true Poisson expectation. As the randomization test *p*-value, we report the proportion of null IOD estimates that were larger magnitude than the observed value.

### Species assortment and co-occurrence

Social network assortment quantifies the extent to which individuals are associated with others that have similar values for a given trait (Newman 2003; James 2015). To determine whether the turtles assort with conspecifics, we used the R package ‘*igraph’* (Csárdi et al. 2024) to build a social network, in which the associations (edges) represented individuals co-occurring in the same capture event. We calculated assortment based on species identity using the ‘*assortnet’* package (Farine 2023). The assortment coefficient can range from –1 to 1; in this case, positive values would indicate that conspecifics were often found together, and negative values would indicate that individuals were often found with heterospecifics (Newman 2003). The data used to calculate species assortment included only focal individuals that had at least one association with another turtle (615 individual turtles that had 1,324 associations). In addition, we also calculated the number of unique individuals from each species that was ever captured with each of the three focal species types (i.e., number of Blanding’s turtles ever caught with a conspecific; painted turtles with a conspecific; snapping turtles with a conspecific; and, number of individuals in each species ever caught with each of its heterospecifics). We used these counts of individuals as our measures of conspecific and heterospecific co-occurrence frequencies.

To compare species assortment and co-occurrence frequencies with null expectations, we simulated several different non-social null models. These null models retained the same number of captures and individuals with the same level of clustering, but permuted the data to generate random co-occurrence (i.e., co-occurrence was independent of turtles’ species, sex, or size) by swapping turtle IDs among capture events within the continuous marsh habitat. We considered four different non-social null model scenarios that differed in the level of spatial and temporal constraint, with further details below.

For the first two null models (models 1 and 2), we used a 3 km spatial constraint, which we expect to be realistic given typical distances travelled by individual turtles at our study site and typical home ranges established in previous studies of our focal species (see Table S2 for details). To construct null model 1, each captured turtle was swapped with another, chosen at random from capture events within 3 km of the trap location, in any year. This process was repeated until all captured turtles had either been permuted or had no remaining candidates that met the spatial constraint to perform additional swaps. In the second case (null model 2), we added a conservative 1-year temporal constraint (i.e., turtles were permuted to other capture events within 3 km of their original capture event location, in the same year).

An important limitation of null models 1 and 2 is that moving turtles between trap locations could disrupt hypothetical species differences in habitat or resource use within the wetland. Furthermore, although other data from the site indicated that trapping sites were used similarly by all three species, we acknowledge that differences in habitat use could generate patterns of positive species assortment based on the species’ spatial distributions, rather than attraction to or avoidance of other turtles. To evaluate this possibility, we built two further null models that had the maximum spatial constraint by preserving the precise trap location (coordinates) where each turtle was captured. Thus, in null model 3, turtles were permuted to the other capture events at the same spatial coordinate location in any year. In null model 4, turtles were permuted to other capture events at the same spatial coordinate location within +/- 1 year of their actual capture. We include these two highly spatially constrained scenarios as a check on whether our findings could be driven by species differences in habitat use within our study site.

For each of the non-social null models described above, we generated 10,000 permuted datasets and derived null expected distributions and randomization tests for each estimate of interest (e.g., species assortment and all co-occurrence frequencies). If the observed estimate fell to the right of the null distribution, we report the *p*-value as the proportion of null estimates larger than the observed value; if the observed estimate fell to the left of the null distribution, the p-value is the proportion of null estimates smaller than the observed value. When discussing the results, we generalized findings to draw the most conservative conclusion from the four null models (with separate null distributions shown in detail in the supplement). Of the 491 capture events, 10 events (with 20 individuals) were missing location coordinates, and these were omitted from all four of the non-social null models.

### Individual rates of aggregation

To further investigate the processes that can drive species assortment, we determined for each of the three focal species the proportion of individuals that had ever been captured with a conspecific. We also compiled null expected distributions and randomization tests for these estimates from the non-social null models described above.

### Body size and aggregation

To investigate whether variation in body size predicts aggregation, we used generalized linear regression models in base R with the binomial distribution and logit link function. Each focal species was modelled separately. We first tested whether a turtle’s body size (carapace length, CCL; Table S3) predicted the probability that it was captured co-occurring with a conspecific at least once (binary response variable; one model per species). Second, we examined whether a turtle’s size (CCL) predicted the probability that it was only captured alone (never co-occurring with a conspecific or heterospecific). A small number of focal individuals whose CCL was not recorded (7 out of 816) were excluded from these analyses.

### Sex assortment within species

We tested for sex assortment using a social network for each species, following the same method as described above, in which the associations were based on conspecifics co-captured in the same trap. Analyses of sex assortment included only individuals with known sex, and compared sex assortment values with the expectations from the four non-social null models using the randomization framework as described above. To ensure these results were robust to uncertainties around turtle sex, we repeated sex assortment analyses including the individuals with indeterminate sex (assigning them as either indeterminate, all male, or all female).

## Results

### Clustering in feeding aggregations

Nearly half of the capture events (45%) involved a dyad or small group of turtles (Fig. 1D). The IOD for the number of turtles per capture event was 1.68, which was 2-3 times greater than expected if the turtles occurred in traps independently (Fig. 1D). The IOD exceeded all null estimates of clustering (*p* < 0.0001).

### Species assortment and co-occurrence

We observed a strong positive assortment of turtle species (assortment = 0.47; SE = 0.01; Fig. 2A), indicating that turtles at our study site aggregated more frequently with conspecifics than heterospecifics. Species assortment also exceeded all estimates from all of the non-social null models (*p* < 0.0001; Fig. 2C, S1A).

**Fig. 2.**
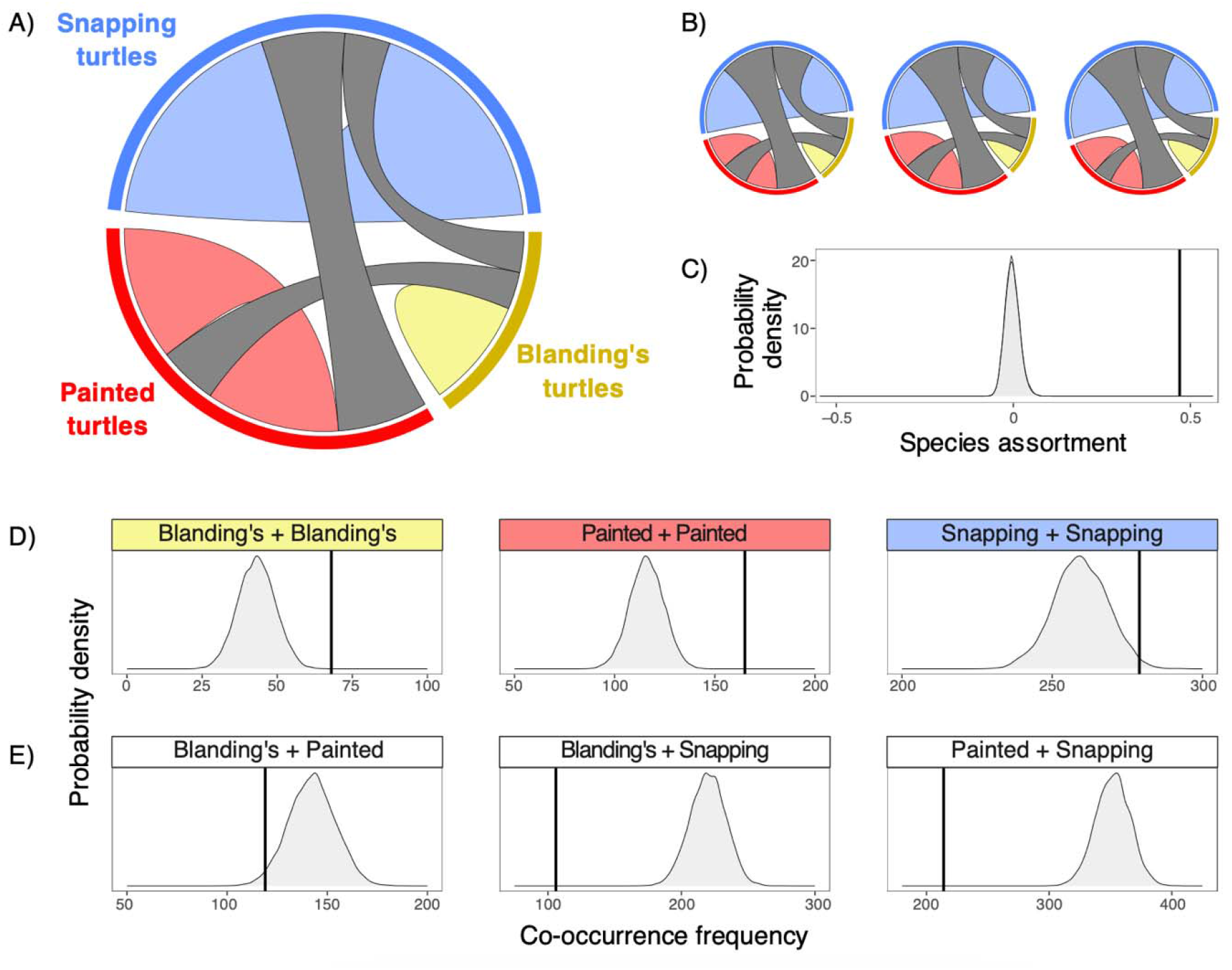
Aggregations of freshwater turtles co-captured in baited hoop traps were positively assorted by species. **(A)** A chord diagram illustrating species assortment. Turtles were often captured with their own conspecifics (coloured arcs), whereas heterospecifics occurred together less often (grey arcs). Arc thickness represents the number of co-captures. **(B)** Examples of the chord diagram derived from iterations of our data for non-social null model 1. In the null datasets, heterospecific turtles generally occurred together at greater frequencies. **(C)** The observed positive species assortment of 0.47 (vertical line) is far greater than the null expectation, which is shown by the shaded probability density. **(D-E)** Co-occurrence frequencies for conspecific and heterospecific turtles. Vertical lines show observed frequencies (number of unique individuals found within each dyad type), and shaded probability density regions show the null expectation for each panel. See Fig. S1 for additional null models

To investigate drivers of species assortment, we examined the frequency of co-occurrence with conspecifics for each species. Both Blanding’s turtles and painted turtles co-occurred with their conspecifics far more often than expected in the non-social null models (Blanding’s turtles: all *p* < 0.0002, and painted turtles: all *p* < 0.0001; Fig. 2D, S1B). Snapping turtles were also captured with other snapping turtles significantly more often than expected (all *p* < 0.02; Fig. 2D, S1B). Examining heterospecific dyads, Blanding’s and snapping turtles co-occurred far less than expected (all *p* < 0.0001), and similarly, painted turtles and snapping turtles also co-occurred far less than expected (all *p* < 0.0001; Fig. 2E, S1C). When considering Blanding’s-painted co-occurrence, these two species co-occurred somewhat less than expected (*p* < 0.03) in the 3km null models (null models 1 and 2). However, we find that Blanding’s and painted turtles co-occurred at a frequency consistent with the null expectation under the maximum spatial constraint (*p* = 0.14 and 0.13 in null models 3 and 4, respectively).

### Individual rates of aggregation

In Blanding’s turtles and painted turtles, 53% and 65% of individuals were co-captured with a conspecific at least once, respectively (Fig. 3A). These individual co-capture rates exceeded the non-social null expectations for all null models (Blanding’s turtles all *p* < 0.001; painted turtles all *p* < 0.0001; see also Fig. S2). In contrast, 65% of the snapping turtles were co-captured with a conspecific at least once, and this fell within the expected range for the 3km null models (all *p* = 0.06; Fig. 3A), and only narrowly exceeded the expectation for null models 3 and 4 (all *p* = 0.04; Fig. S2). Snapping turtles were the most abundantly captured species in our study, so the null expectation for co-capture of two snapping turtles was much higher.

**Fig. 3.**
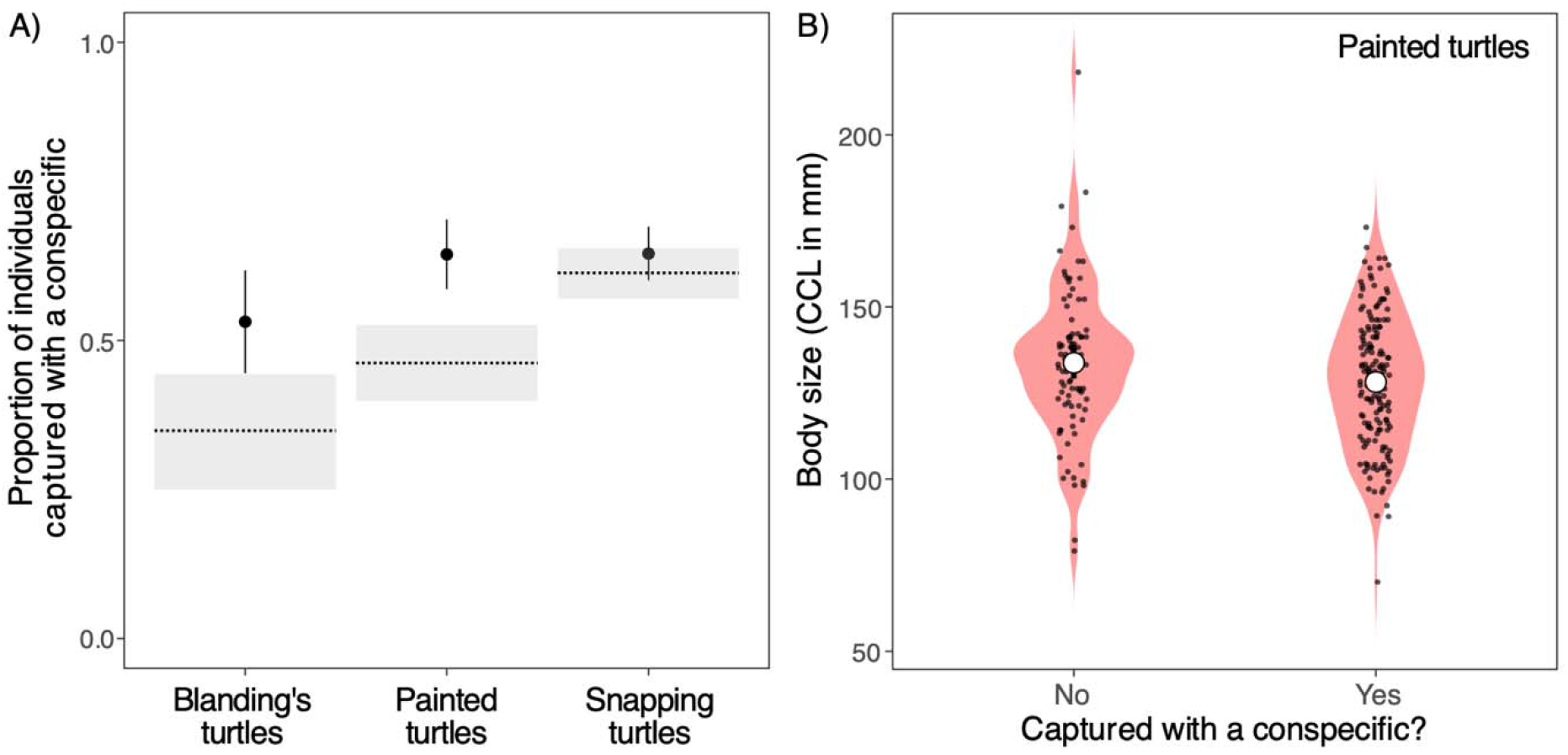
Individual rates of aggregation with conspecifics in freshwater turtles captured in baited hoop traps. A) The proportion of painted turtles and Blanding’s turtles captured with at least one conspecific was greater than expected, whereas the proportion for snapping turtles was within the null expectation. Points show observed values (± 95% CI), and the horizontal dotted lines and grey shaded bands show the null expectation for null model 1 (mean ± 95% percentile region). See Fig. S2 for additional null models. B) Within painted turtles, smaller individuals were slightly more likely to be captured with a conspecific, but this association was not statistically significant when one unusually large painted turtle was excluded. Each small black datapoint represents an individual turtle, and the white circles show means. Note that the black datapoints are jittered on the x-axis for visibility and there is no jitter on the y-axis

### Body size and aggregation

Carapace length did not significantly predict the probability of co-capture with a conspecific in Blanding’s turtles (β = 0.003, SE = 0.006, *p* = 0.620, *n* = 127 individuals) or snapping turtles (β = 0.0001, SE = 0.002, *p* = 0.95, *n* = 429 individuals). We found a weak negative association between carapace length and probability of co-capture with a conspecific in painted turtles (β = – 0.014, SE = 0.007, *p* = 0.04, *n* = 253 individuals; Fig. 3B). However, this association was driven by one particularly large painted turtle (CCL = 218 mm). When this individual was omitted, the association between body size and the probability of conspecific co-capture for painted turtles was not statistically significant (β = –0.012, SE = 0.007, *p* = 0.07, *n* = 252).

When considering the probability of co-capture with any other turtle, carapace length was not a significant predictor in Blanding’s turtles (β = 0.011, SE = 0.007, *p* = 0.09) or snapping turtles (β = –0.001, SE = 0.002, *p* = 0.53). Carapace length was weakly negatively associated with the probability that a painted turtle was captured with any other turtle (β = –0.02, SE = 0.008, *p* = 0.01; Fig. S3), and this association was robust to excluding the particularly large painted turtle (β = –0.02, SE = 0.009, *p* = 0.03).

### Sex assortment within species

All three estimates of sex assortment within species fell within the range of their non-social null expectations (Fig. 4, S4; Blanding’s turtles: *p* = 0.06 to 0.09, painted turtles: *p* = 0.28 to 0.39, snapping turtles: *p* = 0.11 to 0.21). Observed sex assortment estimates were –0.27 for Blanding’s turtles (SE = 0.09, *n* = 59 turtles and 50 edges), –0.05 for painted turtles (SE = 0.03, *n* = 150 turtles and 420 edges), and 0.06 for snapping turtles (SE = 0.04, *n* = 250 turtles and 345 edges; Fig. 4). The three revised sex assortment analyses that included individuals of indeterminate sex, categorized in three different ways, provided the same qualitative results.

**Fig. 4.**
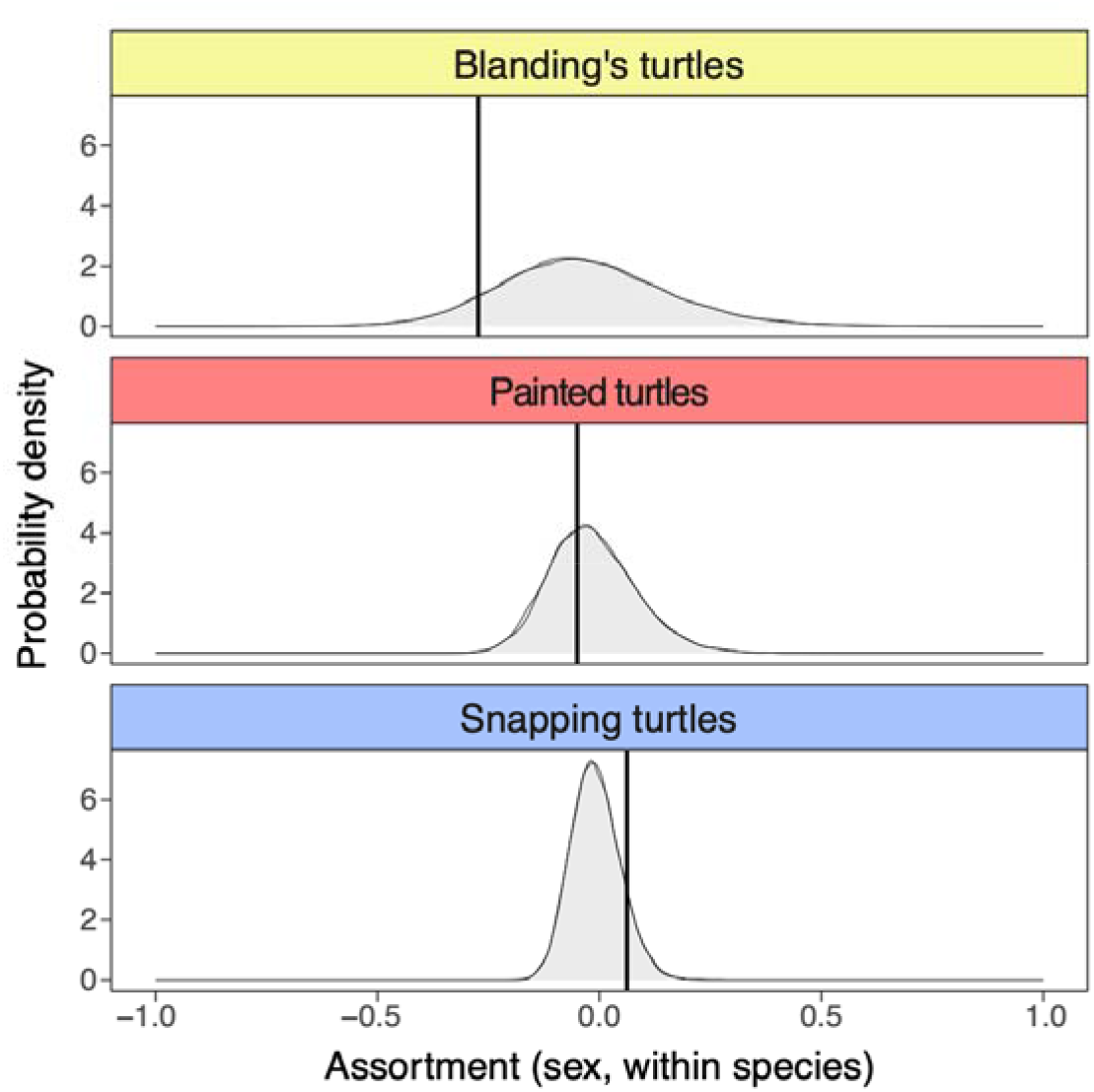
Intraspecific sex assortment in groups of turtles co-captured in baited hoop traps. For each species, the observed sex assortment was within the range expected in non-social null models. Each row shows the estimate for conspecific sex assortment (vertical line), as well as the null expectation shown by shaded probability densities for null models 1 and 2. If the observed assortment (vertical line) is greater than the null predictions (shaded distribution), it would indicate that the frequency of same-sex co-occurrence is greater than expected by random chance. See Fig. S4 for additional null models.

## Discussion

In this study, we used data collected during a long-term population monitoring program to investigate social structure in feeding aggregations of three co-occurring species of freshwater turtle. We found that turtle captures were spatiotemporally clustered, suggesting that the observed aggregations were not random. Feeding aggregations were positively assorted by species, and in particular, Blanding’s and painted turtles were captured with their conspecifics far more often than expected by random chance. Furthermore, while all three heterospecific dyads occurred less often than expected, dyads comprised of snapping turtles and their heterospecifics were especially rare relative to the non-social expectation. Given that maximum carapace length for snapping turtles in our study was 475 mm, compared to 273 mm and 218 mm for Blanding’s and painted turtles, respectively, this result implies that smaller turtles may avoid aggregating with large snapping turtles around potential sources of food, possibly to reduce their predation risk (Williams et al. 2026). The apparent avoidance of snapping turtles by smaller turtles also aligns with observations that snapping turtles are only rarely observed basking with other, smaller turtles (Legler 1956; Obbard and Brooks 1979; Lovich and Gibbons 2021). Interestingly, although painted and Blanding’s turtles often share basking sites (e.g., Lindeman 1999), these species were less willing to aggregate with one another around an underwater food source, suggesting potential competition for resources. To our knowledge, our study is the first to show that species identity drives the composition of freshwater turtle feeding aggregations.

Social network analysis often requires a relatively large number of observations, and our study illustrates how conventional capture data from long-term ecological monitoring programs can be paired with social network analyses to better understand the social behaviour of species that are rare, or difficult to observe directly (see also: Skinner et al. 2023; Clark and Ioannou 2025). The methods we used in this study can be adjusted to study the sociality of any species where individuals can be identified when captured in traps with other individuals, or when monitored sharing space with other individuals (i.e., refuges, hibernacula, or feeding aggregations). Furthermore, these methods are optimal for aquatic environments where direct, underwater observation is challenging (Clark and Ioannou 2025). Although there are some limitations to this approach, which we discuss below, there are also insights to be gained. Adapting existing monitoring datasets, such as the one used in this study, could open many new avenues for the study of the social behaviour of species that are historically underrepresented in behavioural ecology, particularly rare species of conservation concern.

We acknowledge that intraspecific variation in resource selection could shape feeding aggregations differently in more heterogeneous habitats, by altering rates of competition or the probability of interspecific encounters (e.g., Darmon et al. 2012; Corbalán and Debandi 2014). The data for this study were collected in a continuous, large, coastal marsh, across which home ranges of the three focal species overlapped, and previous hand-capture surveys and radio-tracking showed that these species are each able to move through all parts of the study area (Angoh et al. 2019; CMD, unpublished data). Nevertheless, to consider the possibility that habitat use and resource selection could drive our results, we compared observed patterns of assortment and co-occurrence with null models that varied in the level of spatial constraint. In null models 1 and 2, we allowed for individual movement, whereas in null models 3 and.4, we constrained the precise location of each captured turtle to preserve any underlying differences in habitat use. Notably, even the most spatially constrained null models did not generate the strong positive species assortment found in our study (Fig. S1). Thus, the grouping of conspecific turtles during foraging is unlikely to be driven by species differences in habitat use within our study site. Future research could explore how interspecific variation in resource selection among species might further affect the composition of turtle feeding aggregations.

Previous literature suggests that freshwater turtles are attracted to conspecifics (Zipko 1982; Krawchuk et al. 1997; Stollery et al. 2019; Kell et al. 2022; Rouleau et al. 2026). Our results provide further evidence of conspecific attraction and provide insight into potential mechanisms driving turtle feeding aggregations. None of the three species exhibited significant intraspecific sex assortment in within-trap aggregations, so they did not appear to be either searching for or avoiding potential mates while foraging. Nor did turtle body size variation within species strongly predict the probability of aggregation with conspecifics at perceived sources of food. Snapping turtles, the largest species in our study, can exhibit combative and aggressive behaviour with conspecifics (Galbraith et al. 1987; Keevil et al. 2017; Williams et al. 2026). We consequently expected that smaller snapping turtles might avoid interactions with larger individuals. Instead, we found that size did not significantly predict co-capture in snapping turtles; moreover, 40 capture events included two or more male snapping turtles, representing 22% of all capture events in which snapping turtles occurred. Individuals of all three study species are known to have overlapping home ranges, both with their conspecifics and the other focal species (Hammer 1969; Obbard and Brooks 1981; Galbraith et al. 1987; Browne 2003; Rowe and Dalgarn 2010; Angoh et al. 2021). Our study lends support, from a different angle, as to why this home range overlap and aggregation at basking sites may occur, because our findings support a willingness to also aggregate around sources of food.

Social avoidance is also important to consider when describing a species’ sociality (Lusseau et al. 2008; Strickland et al. 2017). This refers to a relative lack of proximity between individuals that share the same environment (Strickland et al. 2017). Hoop traps are not well-suited to characterizing avoidant behaviour because once a turtle enters the trap, it typically remains there until a researcher releases it. The data used in this study do not allow us to determine the order in which the turtles entered the trap, or whether a given individual would have chosen to leave if it could. As such, an important goal for future studies of turtle behaviour is to determine which individuals avoid or displace other individuals, as occurs in species from all non-avian reptile groups (Lang 1987; Firmage and Shine 1996; Lindeman 1999; Ramstad et al. 2012; Itescu et al. 2017; Strickland et al. 2017; Kolanek et al. 2019). Future studies could use camera monitoring, remote PIT tag readers/loggers, GPS transmitters, and/or proximity loggers to determine the order in which individuals enter and leave feeding aggregations. Experiments that “bait” traps with other turtles, rather than food, could also be used to directly test for attraction and avoidance in different turtle species. Studies that remotely monitor tagged individuals may also investigate whether there are consistent individual differences within a species in the tendency to approach, or avoid, conspecifics (Gartland et al. 2021).

In behavioural studies based on passive trapping and mark-recapture data, we note that variation in individual personality can represent a potential confounding factor. In particular, individual personality may affect the likelihood of an animal entering a trap, and it is important to consider that the population sampled from traps may not be representative of the population as a whole (Carter et al 2012; Tesche and Hodeges 2015; Michelangeli et al. 2016; Jolly et al. 2019). Baited traps take advantage of each individual’s need to forage and feed, removing some of this variation. Nevertheless, the different probabilities of trapping “trap-happy’ and “trap-shy” individuals is a well-established issue in mark-recapture studies (Santicchia et al. 2021; Choquette et al. 2024; Cachelou et al. 2026), and individual variation in trapping probability is also well-known in freshwater turtles (e.g., Haydt et al. 2022; Hollender et al. 2022). While we acknowledge this additional source of variation, our focus here was on whether trapped aggregations of turtles showed non-random associations, not on variation in individual capture probability.

The context in which behaviour is observed is particularly important for understanding freshwater turtle sociality. Freshwater turtles partake in a range of behaviours both above and below the water’s surface. Basking is conspicuous and relatively well-studied (e.g., Russell 2016; McKnight et al. 2023; Chessman 2024; Rouleau et al. 2026; McKnight et al 2026), and we built on studies of basking behaviour to characterize a new aspect of freshwater turtle sociality, as it occurs in underwater feeding aggregations. Turtles aggregating around a perceived food source likely have different priorities than when basking, as competition for food may have greater consequences in areas with limited resources than occasional displacement from a basking structure (Polo-Cavia et al. 2011; Pearson et al. 2015). Nevertheless, we found that turtles often formed feeding aggregations with conspecifics and heterospecifics, suggesting that the potential benefits of aggregation (for example, increased vigilance against predators) may outweigh the potential risks of encounter with other turtles. More broadly, our results suggest that there may be a social aspect to turtle foraging itself (i.e., information transfer among individuals), with further research required to test this hypothesis. For species such as snapping turtles that do not commonly bask (Obbard and Brooks 1979; Lovich and Gibbons 2021), quantifying underwater interactions is critical to understanding their social behaviour.

## DATA AVAILABILITY

Data and reproducible code are available at https://doi.org/10.6084/m9.figshare.31934292

## AUTHOR CONTRIBUTIONS

CMD, CM, RD, RJ and JLR designed the study; CM, RJ, and CMD collected the data; CM and RD analyzed the data; CM, CMD, and RD wrote the manuscript, and all authors edited and approved the manuscript.

## ACKNOWLEDGEMENTS

Staff at Ontario Parks generously provided logistical support; thanks especially to E. Slavik, P. Manarome, L. Penner, and J. Matthews. We are grateful to field teams on the Wetlands and Reptiles Project who assisted in collecting the data used in this study, including S.Y.J Angoh, P. Bernardo, L. Browning, P. Catling, J. Choquette, J. Chu, L.C. Comsa, S. Coombes, R. Dillon, M. Fenech, I. Fortin, C. Frenette-Ling, A. Hjort, S. Hudson, J. Larkin, A. Leifso, M. Levac, J.E. Paterson, G. Prince-Badke, K. Ritchie, J. Skuza, B. Talarico, D. Storisteanu, E. Trendos, A. Vanderpas, X. Wang, and A. Whitear. Feedback from two anonymous reviewers greatly improved our manuscript, as did feedback from T. Sherratt, V. Careau, and C. Cullingham on an earlier version. Finally, we gratefully acknowledge the historic and ongoing contributions of First Nations in stewardship of the land and wildlife around the Laurentian Great Lakes, including this study site.

## Statements and Declarations

### ETHICS APPROVAL

All applicable international, national, and/or institutional guidelines for the use of animals were followed. Field methods were approved by the Animal Care Committees of the Royal Ontario Museum (authorizations ROM2008-11, ROM2009-02, ROM2009-21, ROM2010-14, and ROM2011-18), Trent University (authorization 24900), Carleton University (authorization 117299), and the Ontario Ministry of Natural Resources and Forestry (OMNRF authorizations 249 and 291). Additional research authorization was provided by Ontario Parks and the OMNRF (1045769, 1049600,1062210, 1067079) and permits under Ontario’s Endangered Species Act (SR-B-001-10; AY-B-013-11).

### USE OF AI

We made no use of generative artificial intelligence tools in this study.

