## supplement for "A look inside the net: freshwater turtles assort with conspecifics in feeding aggregations"

**Table S1. Trap effort at a coastal marsh on the north shore of Lake Erie during a long-term research and monitoring program (2009-2022)**. Trapping targeted snapping turtles (*Chelydra serpentina),* midland painted turtles (*Chrysemys picta marginata),* and Blanding’s turtles (*Emydoidea blandingii*). Trapping methods were not designed for the current study, so trap effort and locations varied across the study period.

| **Year** | **Number of unique hoop traps set** | **Number of unique trap locations** | **Number of trapping days*** | **Estimated total trap hours**** | **Trapping date range** |
| --- | --- | --- | --- | --- | --- |
| 2009 | 3 | 3 | 3 | 216 | June 23 – 25 |
| 2010 | 3 | 3 | 1 | 72 | July 9 |
| 2011 | 5 | 5 | 23 | 2,760 | May 1 – July 6 |
| 2012 | 5 | 14 | 14 | 1,680 | April 25 – May 19 |
| 2013 | 5 | 7 | 26 | 3,120 | April 28 – June 3 |
| 2014 | 7 | 7 | 15 | 2,520 | May 14 – June 5 |
| 2015 | 7 | 7 | 15 | 2,520 | May 18 – June 5 |
| 2016 | 7 | 9 | 35 | 5,880 | April 26 – June 18 |
| 2017 | 7 | 9 | 15 | 2,520 | May 23 – June 12 |
| 2018 | 7 | 7 | 16 | 2,688 | June 1 – July 6 |
| 2019 | 7 | 7 | 45 | 7,560 | May 21 – August 7 |
| 2020 | *Monitoring program paused due to COVID-19 pandemic* | | | | |
| 2021 | *Monitoring program paused due to COVID-19pandemic* | | | | |
| 2022 | 7 | 7 | 15 | 2,520 | May 16 – 31 |

* *Traps were pulled during inclement weather, or to focus on other project activities such as nesting surveys or radio-telemetry. Therefore, the number of ~24-hour trapping days was sometimes fewer than the full date range of trapping activities.*

** *Trap hours were not reliably recorded in some years. To provide comparable estimates among years, trap hours were estimated based on [number of traps set x the number of 24-hour trapping days x 24 hours].*

**Table S2. A summary of species-specific movement and home range information for the study species, used to inform the spatial constraint employed in the non-social null models.** This includes the maximum straight-line distance that individual turtles were observed moving between at least two capture events during our study period (max. distance moved; mean ± standard deviation, min. – max.), as well as home range and movement information reported in the scientific literature. Straight-line distances of the turtles at our study site during our study period were calculated using the ‘*distHaversine’* function from the ‘*geosphere*’ package in R (Hijmans et al. 2022).

| **Species**  **(n = number of individuals captured at least twice)** | **Max. distance moved (m)** | **Previously reported home range sizes** | **Additional information considered** |
| --- | --- | --- | --- |
| Snapping turtle (*Chelydra serpentina)*  n = 58 | 951 ± 920;  45 – 2762 | 3.4 – 35 ha (Ernst & Lovich 2009; Paterson et al. 2012) | Snapping turtles in Canada typically have larger home ranges than those in the United States of America (Ernst & Lovich 2009). Snapping turtles frequently move overland between wetland habitats (Lovich & Gibbons 2021). |
| Midland painted turtle  (*Chrysemys picta marginata)*  n = 31 | 854 ± 833;  153 –2423 | 89 ha (Jaeger & Cobb 2012)  Home ranges can be several kilometres in length (Browne 2003; Rowe 2003; Rowe & Dalgarn 2010) | Painted turtles are capable of relatively long-distance movement, including often travelling overland from one body of water to another (MacCulloch & Secoy 1983; Ernst & Lovich 2009; Roth & Krochmal 2015). |
| Blanding’s turtle  (*Emydoidea blandingii)*  n = 25 | 445 ± 309;  45 – 1015 | 12 – 60 ha (Angoh et al. 2021)  Home ranges can be several kilometers in length (Fortin et al. 2012; Christensen 2013; Baxter-Gilbert 2014) | Blanding’s turtles are capable of relatively long-distance movement, including often travelling overland to make use of multiple wetlands during their active season (Ernst & Lovich 2009; Fortin et al. 2012; Christensen 2013), but do not always move long distances (Angoh et al. 2021). |

**Table S3. Descriptive statistics for curved carapace length (mm), a measure of body size, for three focal species sampled in this study.**

| **Species** | **Mean** | **SD** | **Range** |
| --- | --- | --- | --- |
| Blanding’s turtle (*Emydoidea blandingii)* | 223.7 | 28.4 | 113 – 273 |
| Midland painted turtle (*Chrysemys picta marginata)* | 130.3 | 19.8 | 70 – 218 |
| Snapping turtle (*Chelydra serpentina*) | 295.8 | 52.1 | 112 – 425 |

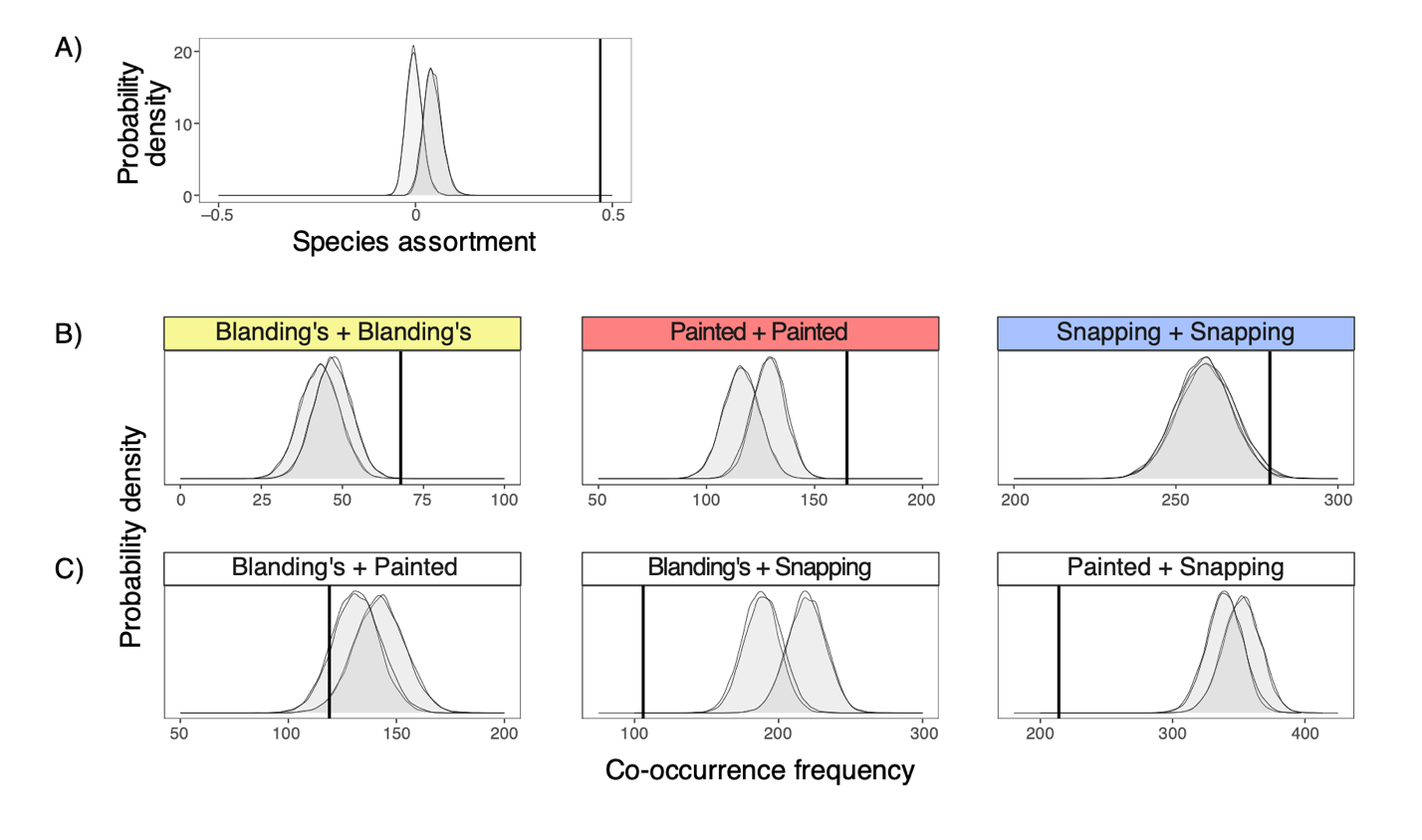

**Figure S1. Aggregations of freshwater turtles co-captured in baited hoop traps were positively assorted by species.** **(A)** The observed positive species assortment of 0.47 (vertical line) is far greater than the null expectation (grey) from four different null model scenarios. Note that four null models are shown here, with extensive overlap among model 1 and 2, and model 3 and 4. **(B-C)** Co-occurrence frequencies for conspecific and heterospecific turtles. Vertical lines show observed frequencies, and the grey shaded regions show null expected distributions for four different null models, as above. The conclusions are broadly unchanged when considering the different null model scenarios.

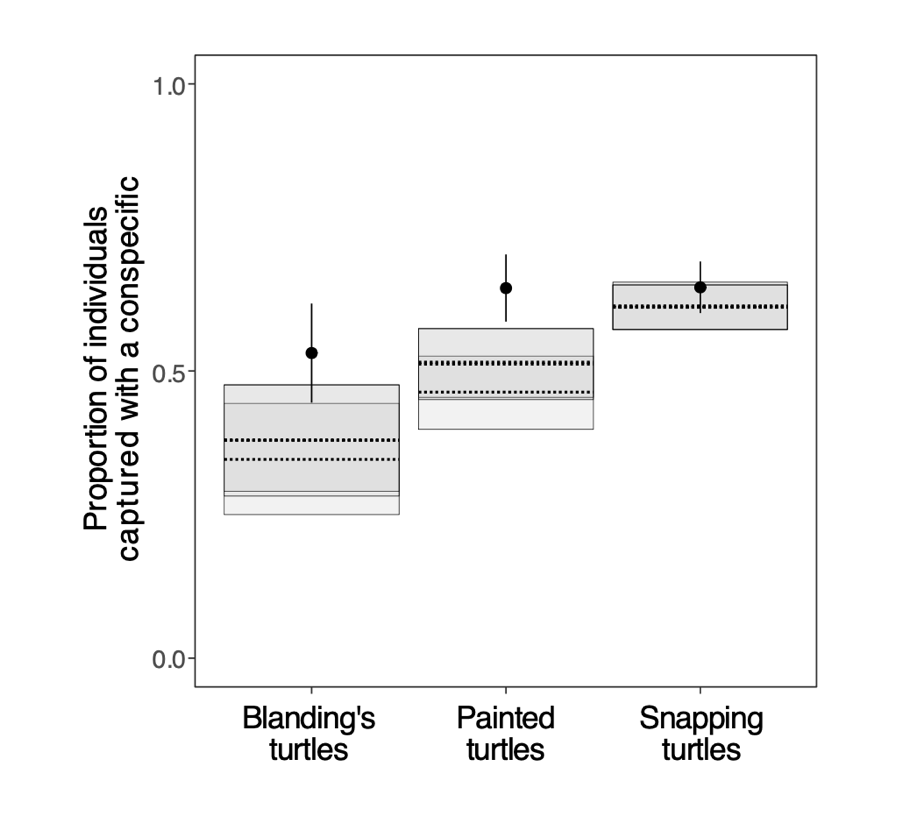

**Figure S2. Individual rates of aggregation with conspecifics in freshwater turtles captured in baited hoop traps.** Points show observed values (± 95% CI), and the horizontal dotted lines and grey shaded bands show four different null model expectations for each species (dotted mean ± grey 95% percentile region). The conclusions are unchanged when considering the four different null model scenarios.

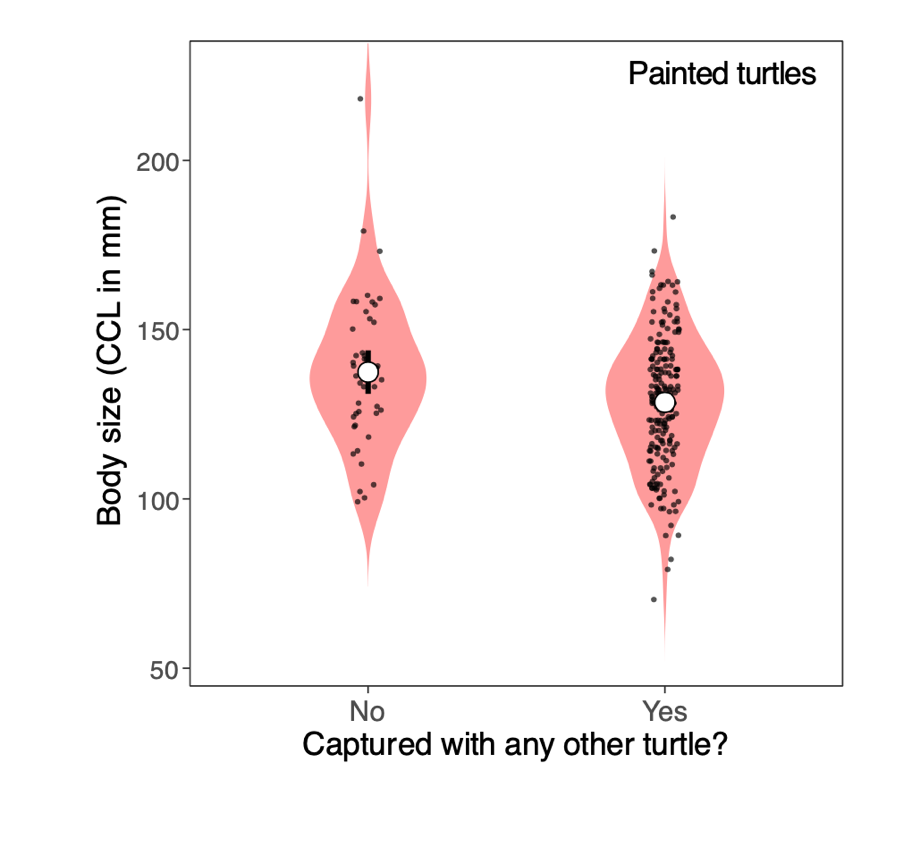

**Figure S3. Within painted turtles, smaller individuals were slightly more likely to be captured with any other turtle (i.e., any species).** Each black datapoint represents an individual painted turtle. The white circles show means and the black bars show ± 95% confidence intervals of the mean for each sample. Note that the 95% confidence interval of the mean for the “Yes” sample falls within the white circle. The black datapoints are only jittered on the x-axis; there is no jitter on y.

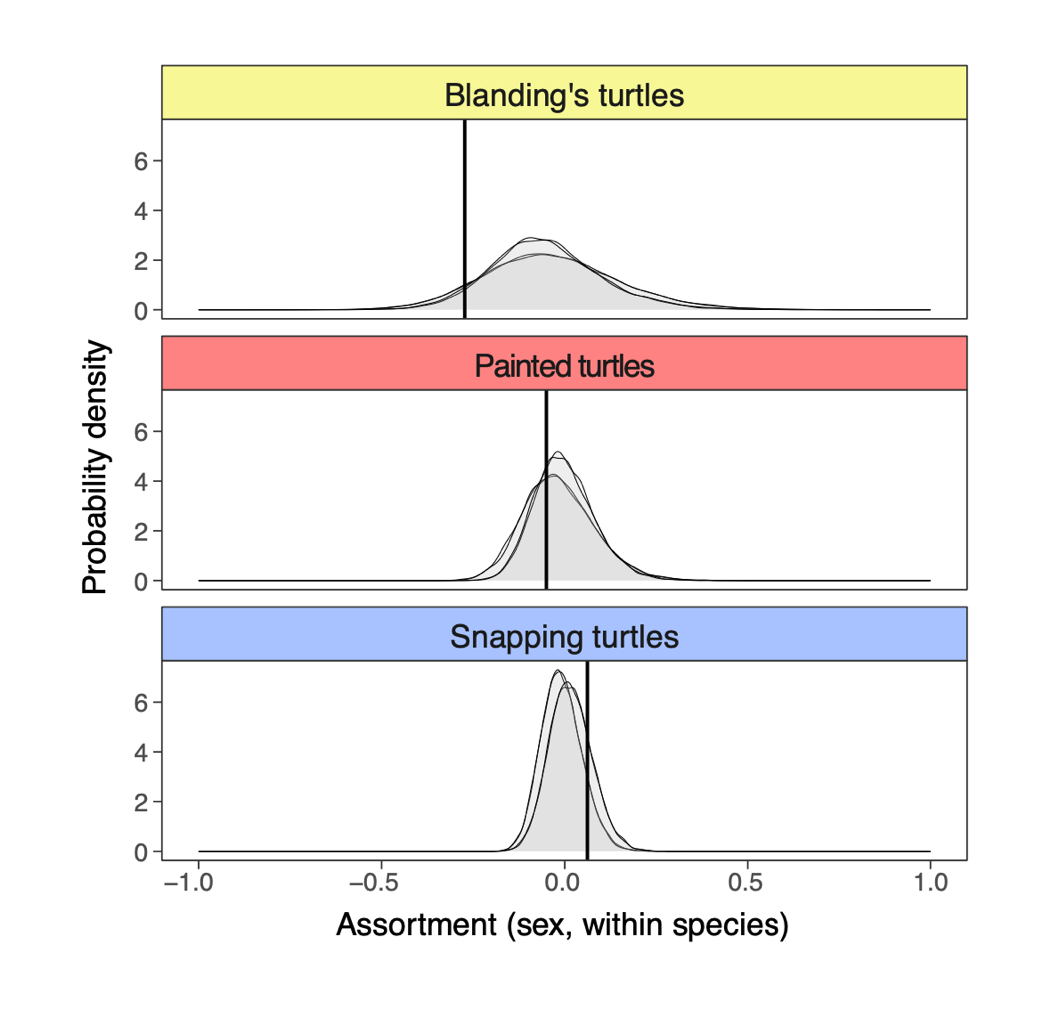

**Figure S4. Intraspecific sex assortment in groups of turtles co-captured in baited hoop traps.** Each row shows the estimate for conspecific sex assortment (vertical line), as well as four different null expected distributions in grey. In all cases, the conclusions are unchanged when considering the four different null model scenarios.
